# Lipid nanoparticle-mediated CRISPR/Cas9 delivery enables efficient trabecular meshwork gene editing in mice

**DOI:** 10.1101/2025.05.17.654623

**Authors:** Yifan Huang, Silin Pang, Linxian Li, Chi Wai Do, Qian Luo, Zongli Zheng, Wenjun Xiong

## Abstract

Lipid nanoparticles (LNPs) have emerged as a transformative platform for mRNA delivery, enabling vaccines and gene editing with transient expression and high cargo capacity. However, their potential for ocular gene editing remains underexplored. In this study, we assessed the transduction efficiency, inflammatory response, and gene editing capability of LNP-encapsulated mRNA in murine eyes. Intravitreal delivery of LNPs achieved targeted mRNA expression in the trabecular meshwork (TM) with superior specificity and efficiency compared to adenoviral or adeno-associated viral vectors, while inducing minimal microglial activation in the retina. Using LNPs co-encapsulating SpCas9 mRNA and sgRNA, we demonstrated efficient CRISPR-mediated knockout (KO) of *Matrix Gla Protein* (*Mgp*), a key inhibitor of TM calcification. *Mgp*-KO eyes exhibited sustained intraocular pressure (IOP) elevation and anterior chamber deepening with normal anterior chamber angle, recapitulating key features of primary open-angle glaucoma (POAG). Chronic IOP elevation led to reactive Müller gliosis and ganglion cell complex thinning, reflecting retinal stress and progressive neurodegeneration. Our findings establish LNP-CRISPR as a safe and efficient system for TM-targeted gene editing, with broad applicability in glaucoma pathogenesis modelling and therapeutic discovery.

## Introduction

Gene therapy has transformed the treatment of retinal diseases, exemplified by the FDA-approved adeno-associated virus (AAV)-based therapy *Luxturna* for Leber congenital amaurosis (*1*). *Luxturna* administers the AAV vector directly into the subretinal space, establishing a new benchmark in retinal gene therapy using viral vectors. In parallel, researchers developed AAV-based therapeutics using intravitreal delivery approaches, which have demonstrated encouraging outcomes in clinical trials targeting conditions like Leber hereditary optic neuropathy and X-linked retinoschisis (*2, 3*). Beyond gene replacement, AAVs have enabled CRISPR-based therapies, such as *EDIT-101* for CEP290-associated LCA (*4, 5*). Despite these breakthroughs, viral vectors-based therapies face many challenges including immune activation risks and difficulties for repeat administration (*6–8*). Additionally, AAVs face limitations such as limited capacity, concerns about vector genome integration at CRISPR-induced DNA breaks, and potential off-target effect when delivering CRISPR/Cas system (*9–11*). These challenges hinder their broader application in gene editing therapies. These limitations highlight the urgent need for innovative delivery systems to overcome current barriers and unlock the full potential of gene therapy for treating retinal degeneration.

Nanoparticle-based platforms offer a promising alternative to viral vectors in gene therapy, combining modularity, scalability, and the ability to deliver larger cargo sizes. These advantages make non-viral nanoparticles versatile delivery systems for diverse applications. A major milestone was the FDA approval of *Onpattro*, an LNP-formulated siRNA therapy for transthyretin amyloidosis (*12*), followed by the rapid development of LNP-mRNA vaccines against SARS-CoV-2 (*13, 14*). LNPs have since demonstrated significant clinical potential, achieving an 87% reduction in transthyretin levels in amyloidosis patients (*15*) and enabling *in vivo* base editing of *PCSK9* in non-human primates (*16, 17*).

Recent studies have investigated the infectivity and therapeutic potential of LNP-mRNA vector in the ocular system. Subretinal injection of various LNP formulations in mice has enabled mRNA delivery to the posterior eye, with expression predominantly observed in non-neuronal cell types, such as the retinal pigment epithelium (RPE) and Müller glia (MG) (*18*). To target retinal neurons, peptide-guided LNPs and thiophene lipids-derived LNPs have been developed, enabling photoreceptor cell transfection in both rodent and non-human primate retinas (*19, 20*). Further optimization of LNP formulations has allowed the delivery of CRISPR/Cas9, base editors, and prime editors, enabling gene editing in the RPE and partial restoration of visual physiology in the *rd12* mouse model (*21, 22*). However, challenges remain, including inflammatory responses and limited transfection efficiency in retinal neurons (*23*). Additionally, the safety and efficacy of LNP vectors in non-retinal ocular tissues remain understudied.

In this study, we evaluated the infectivity and therapeutic potential of LNP-mRNA in the anterior eye segment, demonstrating its efficacy for gene delivery and editing *in vitro* and *in vivo*. Using three distinct ionizable lipids (MC3, ALC0315, SM102) to formulate GFP mRNA-loaded LNPs, we found that SM102-LNPs demonstrated superior transfection efficiency across multiple cell lines while maintaining minimal cytotoxicity. Following intravitreal or intracameral administration in vivo, SM102-LNPs selectively transduced trabecular meshwork (TM) cells with greater specificity and efficiency than AAV vectors, while inducing less retinal microglial activation compared to adenoviral vectors. For CRISPR applications, LNPs co-encapsulated SpCas9 mRNA and sgRNA achieved high editing efficiency in *Rosa26^LSL-tdTomato^* mice, activating tdTomato expression in TM cells. Targeting the gene encoding *Matrix Gla protein* (*Mgp*), which is a critical TM calcification inhibitor, achieved > 60% mRNA knockdown assessed by qPCR or RNA *in situ* hybridization. *Mgp* knockout (KO) mice developed chronic ocular hypertension, exhibiting sustained elevated intraocular pressure (IOP), anterior chamber elongation, and ganglion cell complexes (GCCs) thinning. Together, these results establish SM102-LNPs as an efficient and well-tolerated platform for TM-targeted gene delivery and CRISPR-based genome editing, providing a valuable tool for glaucoma research and therapeutic development.

## Results

### SM102-derived LNP enables strong GFP mRNA expression in cultured ocular cell lines

The three FDA-approved LNP-based therapeutics (Onpattro, Spikevax, and Comirnaty) each employ distinct ionizable lipids: DLin-MC3-DMA (MC3), ALC-0315, and SM-102, respectively. Capitalizing on their proven clinical safety profiles and efficacy, we selected these lipids to formulate three GFP mRNA-loaded LNPs (MC3-GFP, ALC0315-GFP, and SM102-GFP) using identical molar ratios of other elements (Fig. 1A). All three LNPs showed narrow size distribution, with a hydrodynamic diameter ranged from 120 to 160 nm, a polydispersity index (PDI) value < 0.1, positive zeta-potentials and encapsulation efficiencies > 90% (Fig. 1B-E). Among these, SM102-GFP had the smallest size (∼ 120 nm) and highest zeta-potential (> 15 mV).

**Figure 1.**
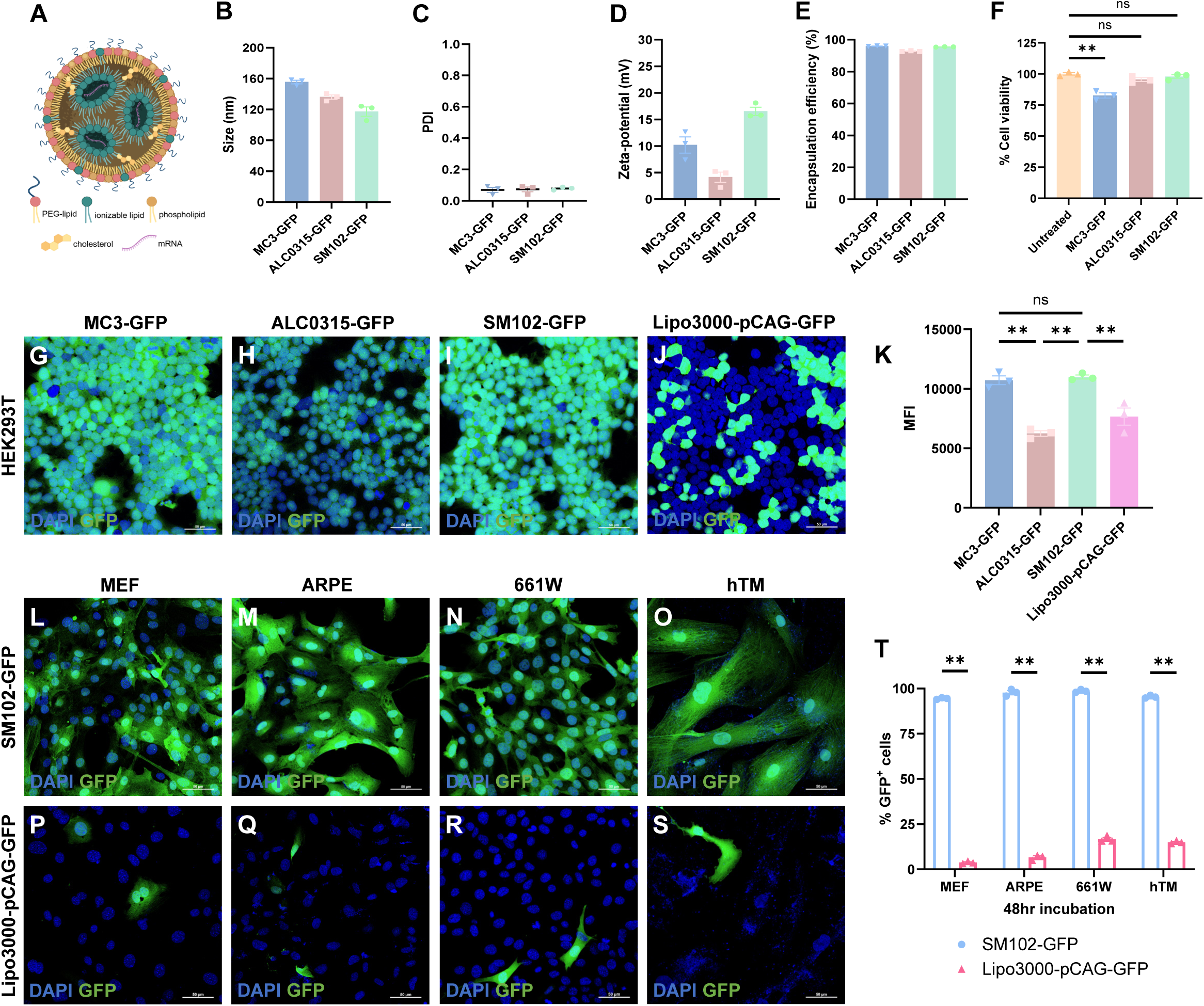
SM102-derived LNP enables strong GFP mRNA delivery in cultured ocular cell lines. (A) Schematic illustration of LNP-mRNA formulation and structure. (B-E) Characterizations of MC3-GFP, ALC0315-GFP and SM102-GFP, including hydrodynamic diameter (B), polydispersity index (PDI) (C), zeta-potential (D), and encapsulation efficiency (E). (F) Cell viability assessed by MTT assay following 24hr treatment with different LNP formulations. (G-J) Representative images showing GFP expression induced by different ionizable lipids-based LNPs and Lipo3000-pCAG-GFP in HEK293T cells. (K) Mean fluorescence intensity (MFI) of GFP signals induced by different LNPs and Lipo3000-pCAG-GFP in HEK293T cells. (L-S) Representative images showing GFP expression after 24hr incubation of SM102-GFP (L-O) or Lip3000-pCAG-GFP (P-S) in different cell lines including MEF, ARPE, 661W and hTM. (T) Quantification of GFP-positive cell numbers in different cell lines at 48hr post-treatment. Data shown as Mean ± SEM, n = 3 per group. **P < 0.01; ns, not significant difference by one-way ANOVA analysis with Tukey test (F, K) and two-tail unpaired student t-test (T). Scale bars: 50 μm.

We compared the mRNA delivery efficiency of the three LNP variants in HEK293T cells, and SM102-GFP and MC3-GFP enabled stronger GFP expression than ALC0315-GFP (Fig. 1G-I, 1K). The relatively higher positive surface charge of SM102-GFP and MC3-GFP may facilitate their cellular uptake through electrostatic interaction with cell membranes (Fig. 1D). To examine the potential cell toxicity of LNPs, 3-[4,5-dimethylthiazol-2-yl]-2,5 diphenyl tetrazolium bromide (MTT) assay was performed, and the results showed that cells treated with SM102-GFP and ALC0315-GFP had comparable viability to untreated cells, while MC3-GFP-treated cells had significantly decreased cell metabolic activity, implying that MC3-derived LNPs might have higher cellular toxicity (Fig. 1F). To sum up, SM102-based LNP outperforms the other two LNPs, with higher mRNA delivery efficiency and lower cell toxicity.

SM102 ionizable lipid-based LNP-mRNA formulation was further tested in multiple cultured cell lines, including mouse embryonic fibroblast (MEF), mouse photoreceptor cell line (661W), human retinal pigment epithelium cell line (ARPE), and human primary trabecular meshwork cell line (hTM). At 1 ng/μL mRNA dose, SM102-GFP achieved remarkable >95% transfection efficiency in all tested cell lines (Fig. 1L-O, T), dramatically outperforming Lipofectamine 3000-delivered plasmid DNA (Fig. 1P-T). These findings demonstrate that LNP-mRNA systems can effectively overcome the transfection barriers typically encountered in ocular cell models, highlighting their significant potential for ophthalmic research and therapeutic applications.

### Intravitreal delivery of LNP enables targeted mRNA expression in the trabecular meshwork (TM)

Building on our *in vitro* success, we evaluated SM102-GFP delivery in mouse eyes via intravitreal injection (Fig. 2A), comparing its performance to AAV2 (AAV2-CMV-GFP) and Ad5 adenovirus (Ad-CMV-GFP). Immunohistochemical analysis one week post-injection revealed distinct tropism patterns: AAV2-CMV-GFP transduced multiple retinal cell types including retinal ganglion cells (RGCs), amacrine cells (ACs), MGs, and optic nerve head (ONH) (Fig. 2B), while Ad-CMV-GFP and SM102-GFP showed limited retinal transduction (Fig. 2C, D). This differential penetration likely reflects physical barriers imposed by the inner limiting membrane (ILM), which restricts access of larger particles to retinal neurons. Notably, both Ad-CMV-GFP and SM102-GFP demonstrated robust GFP expression in the TM, the phagocytic tissue regulating aqueous humor drainage (Fig. 2C, D). GFP-positive cells localized anterior to Schlemm’s canal (SC), which was outlined by collagen-I staining, confirming TM-specific transduction (Fig. 2E-G). Quantification revealed higher TM infectivity for LNP and adenovirus versus AAV (Fig. 2H), with GFP uniformly distributed across outflow facilities in cornea flat-mounts (Fig. 2I).

**Figure 2.**
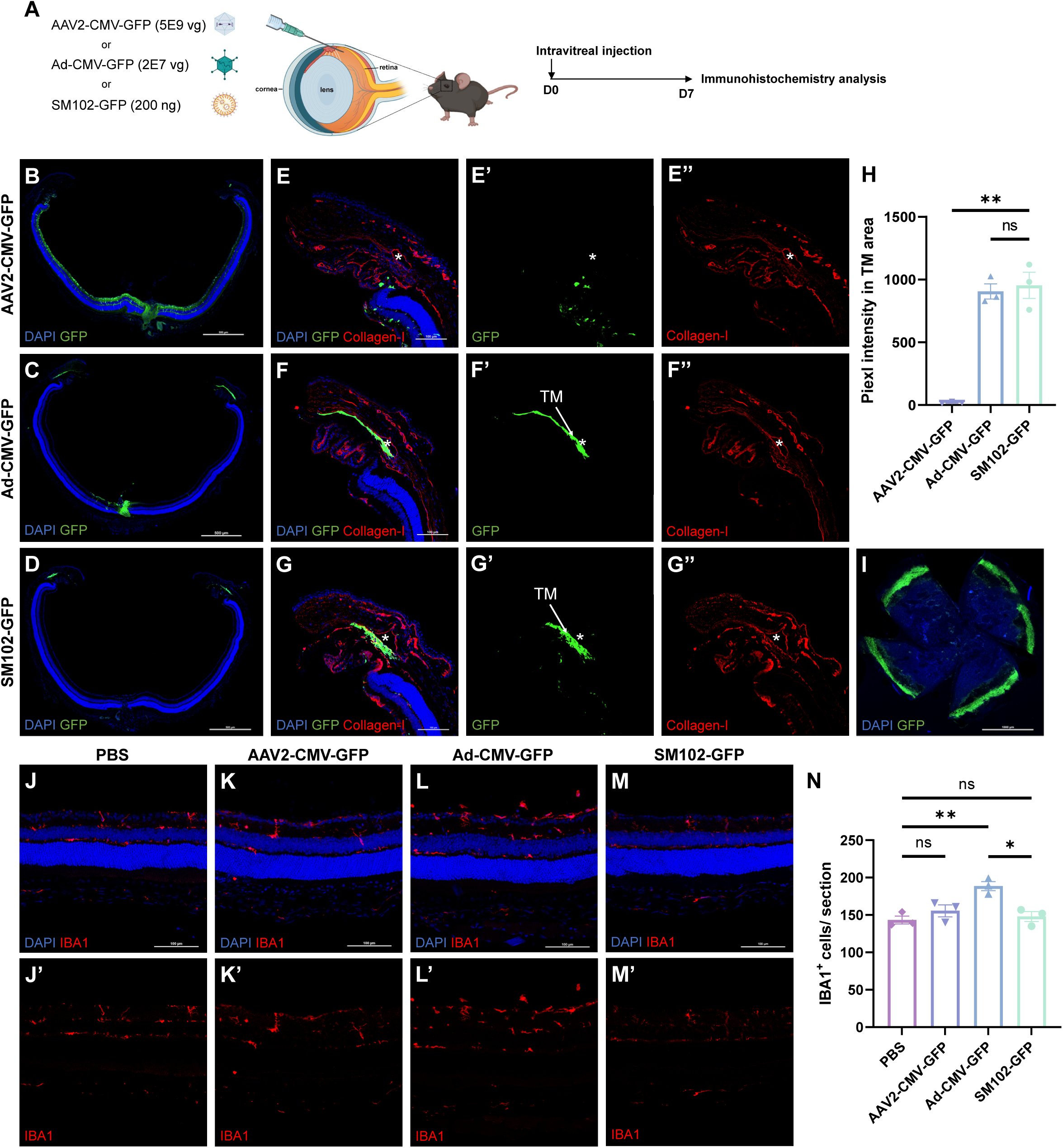
Intravitreal delivery of SM102-GFP LNPs selectively targets trabecular meshwork (TM) in mouse eyes. (A) Experimental workflow for intravitreal injections and tissue processing. (B-G) Representative images of retinal sections showing transduction patterns of AAV2-CMV-GFP, Ad-CMV-GFP, and SM102-GFP. Collagen-I staining (red) delineates Schlemm’s canal (SC, pointed out by *). TM indicates trabecular meshwork. Scale bars: 500 μm for whole retinal sections; 100 μm for zoom-in images. (H) Quantification of MFI in TM areas of AAV2-CMV-GFP, Ad-CMV-GFP or SM102-GFP injected eyes. (I) Anterior segment flat-mount showing pan-ocular GFP expression in outflow tissues. Scale bar: 1000 μm. (J-M) Retinal sections immunostained for IBA1 (microglia marker) following different treatments. Scale bars: 100 μm. (N) Quantification of IBA1-positive cells in retinal sections. Data shown as Mean ± SEM, n = 3 per group. *P < 0.05; **P < 0.01; ns, not significant difference by one-way ANOVA analysis with Tukey test.

Neuroinflammatory responses in the retina are modulated by microglial cells, the resident immune cells in the central nervous system. To investigate the neuroinflammatory profiles of the three gene therapy vectors, we performed anti-IBA1 staining in retinal sections to see whether there was increased microglia activation. The number of IBA1-positive cells was elevated in Ad-CMV-GFP-injected retinas but unchanged in AAV2-CMV-GFP or SM102-GFP-treated eyes versus PBS controls (Fig. 2J-N). These findings position SM102-LNPs as a superior vector for TM targeting, combining high specificity, efficient transduction, and minimal immunogenicity.

Next, we performed a dose study of LNP-mRNA in TMs. The levels of GFP expression and number of TM cells infected showed a dose-dependent correlation across the range of 20 ng to 100 ng of mRNA per injection (Fig. S1B-E), plateauing at 100 ng without further enhancement at 200 ng (Fig. S1A, B, E). Kinetic analysis revealed rapid onset of GFP expression (detectable by 4 hours post-injection), peaking at 24 hours and persisting for one week (Fig. S2A-F). Expression declined by three weeks (Fig. S2G-I), establishing SM102-LNPs as an ideal platform for transient therapeutic applications including gene editing.

### LNP-CRISPR mediates efficient gene editing in TM

Given the potent TM infection achieved by LNP-mRNA, we next investigated whether LNP could efficiently deliver CRISPR/Cas9 system for TM gene editing. The Cre reporter mouse *Rosa26^LSL-tdTomato^* has three repeated stop codon cassettes that hinder tdTomato expression (Fig. 3A). CRISPR-mediated excision of two repeat cassettes was reported to be sufficient to activate the expression of downstream tdTomato (*24*). We generated SM102-SpCas9-sgtdT vector, which co-encapsulated SpCas9 mRNA and a synthesised sgRNA (weight ratio 1:1) with 2’-O-methyl and phosphorothioate modifications targeting the stop cassettes in the *Rosa26^LSL-tdTomato^* genome. SM102-SpCas9-sgtdT showed similar particle size, PDI, zeta potential and mRNA encapsulation efficiency as SM102-GFP (Fig. 1B-E, S3A-D). Co-encapsulated SM102-SpCas9-sgtdT was intravitreally delivered to the *Rosa26^LSL-tdTomato^*mice, and the eyes were harvested for examination one-week post-injection. Retinal section and anterior segment flat-mount images showed SM102-SpCas9-sgtdT enabled robust tdTomato expression in the TMs (Fig. 3B, S3G). tdTomato-positive cells positioned near the SC, without overlap with pigmented ciliary epithelium or RPE (Fig. 3C). α-SMA staining further validated TM-specific editing, showing clear demarcation from ciliary muscle (Fig. 3D). Quantification analysis showed that around 66.1% TM cells were successfully edited with at least one allele of the tdTomato (Fig. 3E, F). Overall, the results indicated that CRISPR/Cas9 system intravitreally-delivered by LNP mediated potent gene editing in the TMs, highlighting the robustness of LNP-CRISPR in TM gene manipulation.

**Figure 3.**
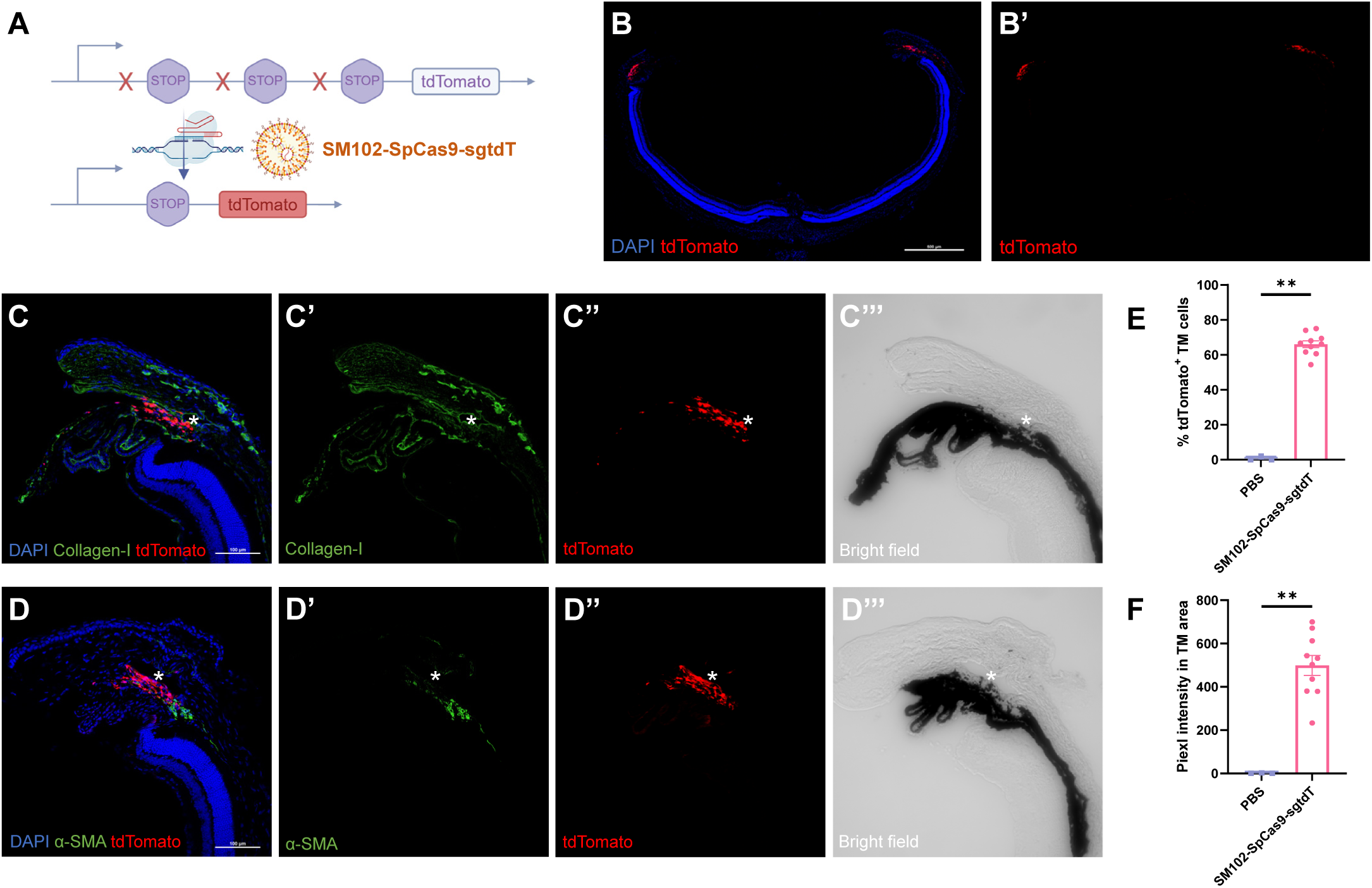
LNP-CRISPR mediates efficient gene editing in mouse TM. (A) Strategy for tdTomato activation in *Rosa26^LSL-tdTomato^* mice. (B) Representative images of retinal sections showing robust and specific tdTomato expression in retinal outflow facility. Scale bars: 500 μm. (C, D) Co-staining with Collagen-I (green) or α-SMA (green) confirms TM-specific editing. * indicates SC. Scale bars: 100 μm. (E, F) Quantification of tdTomato-positive TM cell numbers and MFI. PBS (n=3); SM102-SpCas9-sgtdT (n=10). Data shown as Mean ± SEM. **P < 0.01; ns, not significant difference by two-tail unpaired student t-test.

Intracameral injection is a safe drug delivery route to the anterior segments. Here we also compared the infection patterns of intravitreally and intracamerally delivered SM102-SpCas9-sgtdT in *Rosa26^LSL-tdTomato^* mouse eyes (Fig. S3E). The LNP by both injection routes could transfect TMs without retinal or lens transduction (Fig. S3F, H). The anterior segment flat-mounts by intracameral injection showed strong tdTomato expression in the retinal outflow facilities as well, but we could see some signals in a small portion of the cornea (Fig. S3I). We speculated that intracameral injection could lead to the disruption of corneal epithelium integrity at the site of injection, allowing the epithelium infected by the LNP. Nonetheless, both intravitreal and intracameral injections are viable administration routes for targeting TM by LNP vectors.

### LNP-CRISPR-mediated *Mgp* KO in TM induces chronic ocular hypertension in mice

The TM serves critical functions in draining the aqueous humor and maintaining IOP, and dysfunction of TM is strongly associated with glaucoma pathogenesis (*25–28*). Next, we conducted an *in vivo* study using LNP-CRISPR to knock out specific TM gene for functional analysis. Analysis of published single-cell RNA sequencing database of TMs in mouse species revealed that *Mgp* is one of the most highly expressed genes in mouse TM tissues (*29*). We further verified its expression in TM by qPCR analysis, which showed that *Mgp* had a 134-fold higher expression in TMs than in retinas (Fig. S4A). The MGP protein plays a role in inhibiting calcification and protecting soft tissue from stiffness (*30–33*). It has been shown that *Mgp* conditional KO in TMs causes progressive intraocular hypertension in mice (*34*). We selected *Mgp* as the target gene for establishing a glaucoma model using the LNP-CRISPR system.

We designed two sgRNAs targeting exons 1 and 4 of mouse *Mgp* coding region, which were co-encapsulated with SpCas9 mRNA in LNPs at 1:1:1 weight ratio (Fig. 4A). The SM102-SpCas9-sgMgp vector exhibited similar properties to our previously characterized SM102-GFP and SM102-SpCas9-sgtdT (Fig. 1B-E, S3A-D). Intravitreal injection of SM102-SpCas9-sgMgp (100 ng total RNA/eye) was performed to the right eyes of mice, and the contralateral left eyes were uninjected controls. Another group of mice were injected with SM102-SpCas9-sgtdT in the right eyes to account for any potential effects of the LNP vector, SpCas9 mRNA, or nontarget sgRNA. Mice were sacrificed at 3 weeks- or 15 weeks-post injection for next generation sequencing (NGS), qPCR and RNAscope assay to determine *Mgp* KO efficiency (Fig. 4B). NGS results showed that the two sgRNAs achieved an average of 6.0% and 1.7% editing events in the iridocorneal angle tissues (Fig. 4C, S4B). These values likely underestimated true TM editing efficiency due to contamination from non-TM cells in dissected tissues and the inability to detect larger deletions resulting from end-joining between the two sgRNA target sites. Consistent with this interpretation, qPCR analysis showed more than 60% reduction in *Mgp* mRNA (Fig. 4D), while RNAscope assay quantification showed 72.2% loss of *Mgp* mRNA signals in *Mgp*-KO eyes (Fig. 4E-H). Notably, the mRNA signals of *Myocilin*, another TM gene, remained unchanged, confirming preservation of TM structure and the specificity of our editing approach. These convergent results confirm that our SM102-SpCas9-sgMgp system achieves efficient and specific *Mgp* KO in the TM.

**Figure 4.**
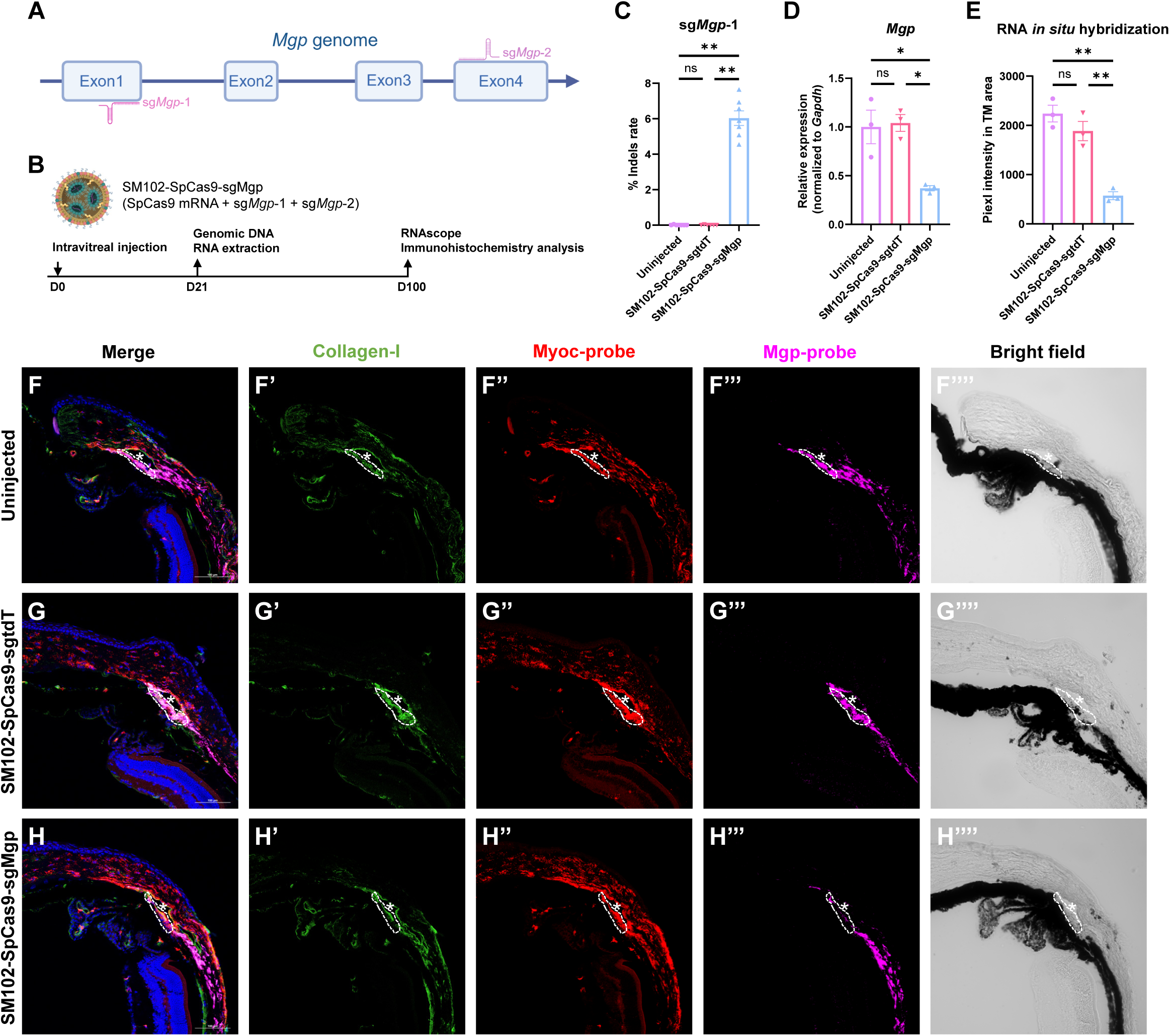
TM-specific *Matrix Gla Protein* (*Mgp*) knockout (KO) using LNP-CRISPR. (A) sgRNA targeting strategy for *Mgp* KO. (B) Experimental timeline. (C) Quantification of insertions and deletions (Indels) at sg*Mgp*-1 targeting site in SM102-SpCas9-sgMgp (n=7) or SM102-SpCas9-sgtdT (n=6) treated or untreated (n=7) TM tissues by using next generation sequencing. (D) qPCR analysis of *Mgp* mRNA expression in SM102-SpCas9-sgMgp or SM102-SpCas9-sgtdT treated or untreated TM tissues (n=3 for all groups, normalized to Gapdh mRNA). (E) Quantification of MFI of Mgp signals in TM areas of SM102-SpCas9-sgMgp or SM102-SpCas9-sgtdT injected or uninjected eyes by using RNAscope assay (n=3 for all groups). (F-H) Representative images of retinal sections with Myocilin and Mgp mRNA *in situ* hybridization showing *Mgp* knockdown in *Mgp*-KO eyes compared to control eyes. * indicates SC. Data shown as Mean ± SEM. *P < 0.05; **P < 0.01; ns, not significant difference by one-way ANOVA analysis with Tukey test. Scale bars: 100 μm.

Longitudinal monitoring revealed progressive ocular changes in *Mgp*-KO eyes exhibiting key features of primary open angle glaucoma (POAG) (Fig. 5A). IOP steadily increased to a plateau of 17.6 mmHg by day 90 that persisted through day 200 (Fig. 5B). Whole-eye optical coherence tomography (OCT) demonstrated increased axial length (AL) and anterior chamber depth (ACD) in *Mgp*-KO eyes compared to controls (Fig. 5C-E, L, M), with anterior segment OCT confirming iridocorneal angle widening and anterior chamber enlargement (Fig. 5F-H). Despite preserved retinal morphology (Fig. 5I-K) and normal photoreceptor and bipolar cell function by electroretinography (ERG) at day 200 (Fig. S4G-L), we observed progressive GCC thinning (Fig. 5N) and GFAP-positive Müller gliosis (Fig. 5O-Q), indicating retinal stress responses to chronic elevated IOP. Notably, immunostaining and pattern-ERG revealed no significant ganglion cell loss by day 200 (Fig. S5), suggesting this model recapitulates the pathophysiology of early-stage POAG while maintaining retinal function. Together, these results establish that LNP-CRISPR-mediated *Mgp* KO in TM tissue creates a robust chronic ocular hypertension model, providing a valuable tool for glaucoma research and therapeutic development.

**Figure 5.**
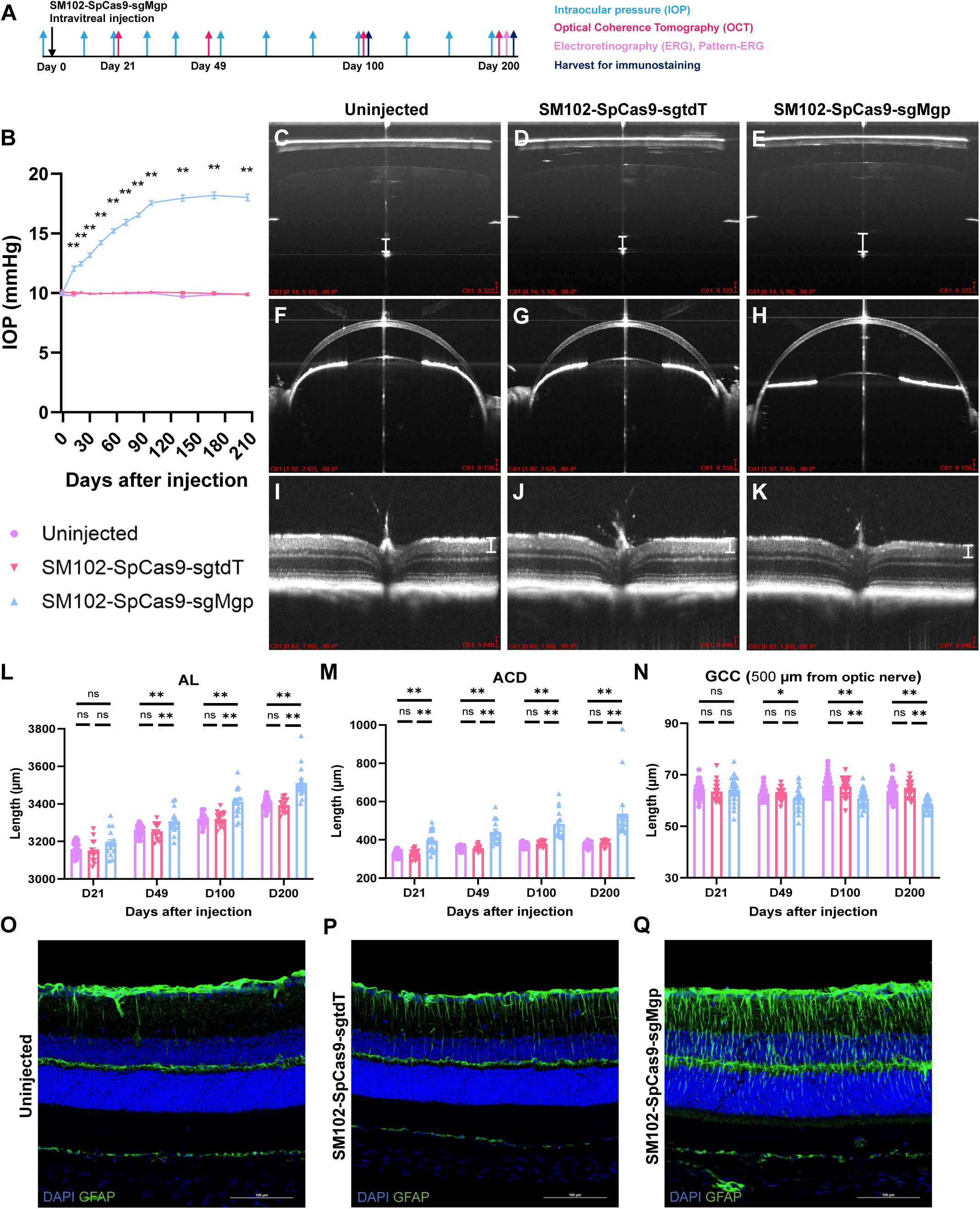
*Mgp* KO induces chronic ocular hypertension and glaucomatous changes. (A) Experimental timeline. (B) IOP results of eyes uninjected (n=20) or intravitreally injected by SM102-SpCas9-sgMgp (n=10) or SM102-SpCas9-sgtdT (n=10). (C-K) Representative OCT images showing elongated axial length (AL) and anterior chamber death (ACD), open iridocorneal angle, normal retinal morphology and thickness of *Mgp*-KO eyes compared to control eyes on Day 200-post injection. Scale bars: 0.322 mm for whole eye images; 0.150 mm for anterior chamber images; 0.048 mm for retinal images. (L-N) Quantification of AL, ACD and ganglion cell complex (GCC) length of uninjected (n=30), control LNP-injected (n=14) and *Mgp*-KO eyes (n=16). (O-Q) GFAP staining showing Müller gliosis in *Mgp*-KO eyes compared to control eyes on Day 100-post injection. Scale bars: 100 μm. Data shown as Mean ± SEM. *P < 0.05; **P < 0.01; ns, not significant difference by one-way ANOVA analysis with Tukey test.

## Discussion

While CRISPR-Cas9 technology represents a transformative approach for treating genetic disorders, current delivery systems require substantial optimization to address efficacy and safety concerns. For example, the first-in-human CRISPR-Cas9 clinical trial using *EDIT-101* is currently paused as only 21.4% participants obtained visual improvements (*35, 36*). In this trial, AAV5 vectors delivered CRISPR components to photoreceptors, raising two critical safety issues: First, the potential for off-target effects remains a significant concern. Unintended genome edits or indels at non-target loci could accumulate due to long-term AAV-mediated transgene expression. Second, studies have revealed that AAV vectors preferentially integrate into CRISPR-Cas9-induced double-strand breaks at rates up to 47% (*9*). This integration phenomenon poses substantial risks, as it could lead to insertional mutagenesis or disrupt normal gene regulation.

An alternative delivery system, LNP-mRNA, provides several advantages over AAV-based delivery system for CRISPR-Cas9 therapeutics. First, multiple studies, including our own work presented here, have proved that LNPs can effectively encapsulate both Cas9 mRNA and sgRNA and realizes gene editing efficiently (*16, 37, 38*). Second, the transient nature of mRNA-mediated expression limits Cas9 activity to a defined window (typically 1-2 weeks, as we observed with GFP expression in TM cells), significantly reducing the risk of cumulative off-target effects compared to persistent AAV-mediated expression. Third, and perhaps most importantly, it eliminates the possibility of genome integration, which is a major safety concern for AAVs. Additionally, the LNP platform allows for repeat dosing to enhance editing efficiency, as evidenced by the clinical success of both mRNA vaccines (e.g., COVID-19 vaccines) and therapeutic siRNA delivery (e.g., *Onpattro*). This flexibility represents a significant advantage over AAV vectors, which are limited by pre-existing immunity and cannot be readministered effectively.

Contrary to expectations based on AAV2 behavior, intravitreally administered LNPs did not efficiently transduce retinal neurons. Instead, we observed robust transduction specifically in the TM. This distinct tropism likely results from three factors: (1) the larger size of LNPs (∼120 nm) preventing penetration through the inner limiting membrane (ILM). This may explain why we did not observe LNP infection in retinas which was reported in other studies using smaller size of LNPs with different formulations (*39, 40*); (2) subsequent aqueous humor circulation directing LNPs to outflow tissues; (3) the phagocytic nature of the TM cells (*41*). This natural targeting mechanism provides inherent specificity for TM transduction without requiring complex surface modifications.

Capitalizing on the TM’s crucial role in IOP regulation, we targeted this tissue to establish a glaucoma model. Here, we chose *Mgp* as the glaucoma induction gene based on its high expression level in TM tissues and its important role for keeping tissue soft by inhibiting calcification. Although the functions of *Mgp* in bone and cardiovascular systems are well-documented (*33, 42*), its importance in TM physiology remains elusive. Using the LNP-CRISPR platform, we achieved substantial *Mgp* knockdown in TM tissue, with qPCR showing >60% reduction in mRNA levels and RNAscope analysis demonstrating 72.2% loss of Mgp signal (Fig. 4D, E). This genetic manipulation resulted in progressive IOP elevation that stabilized at around 18.0 mmHg, along with characteristic anatomical changes of POAG, including anterior chamber deepening with a normal open iridocorneal angles (Fig. 5C-N). Although we did not observe massive RGC loss even at late time point (Fig. S5J), a gradual GCC thinning became evident starting from day 49 post-LNP injection (Fig. 5N). Müller gliosis, as confirmed by increased GFAP staining, was observed in the eyes with *Mgp* knockdown at day 100 but not in the control LNP injected eyes, suggesting that retinal stress resulted from chronic IOP increase (Fig. 5O-Q). The lack of RGC loss is likely due to the moderate IOP increase (Fig. 5B). Reports have shown that the mouse RGCs could accelerate degeneration only if IOP maintained over 20 mmHg for several months (*43–45*). While existing approaches, including microbead occlusion (*46*), laser photocoagulation (*47*), pharmacological induction (*48*), and silicone oil injection (*49*), produce acute IOP elevation and severe RGC loss, our model may better recapitulate chronic glaucoma pathogenesis. To potentially enhance the disease phenotype, future iterations could co-deliver mRNA encoding calcification factors such as BMP2/4 and RUNX2 to further disrupt TM outflow function (*50*).

Beyond disease modeling, this LNP-CRISPR platform shows significant therapeutic potential. The same delivery strategy could be adapted to KO disease-associated genes such as mutant *Myocilin* or to modulate regulators of TM contractility including *ROCK1* and *ROCK2* for lowering IOP (*51–53*). Our findings therefore accomplish two important advances: they demonstrate LNP-mRNA’s unique tropism for ocular outflow tissues, and they establish CRISPR-based genetic manipulation as a powerful approach for both studying and potentially treating glaucoma and other ocular disorders.

## Methods

### Materials

Dlin-MC3-DMA and SM102 were purchased from BioFine International Inc. ALC-0315, 1,2-distearoyl-sn-glycero-3-phosphocholine (18:0 PC, DSPC), and 1,2-dimyristoyl-rac-glycero-3-methoxypolyethylene glycol-2000 (DMG-PEG2k) were purchased from Avanti Polar Lipids. Cholesterol was purchased from Sigma Aldrich. sgRNA including sgtdT, sg*Mgp*-1 and sg*Mgp*-2 were custom synthesized with 2’-O-methyl and phosphorothioate modifications at the first three 5’ and 3’ terminal RNA residues from GenScript. Rabbit anti-Collagen-I (ab34710), rabbit anti-α-SMA (ab124964) and rabbit anti-Rbpms (ab152101) primary antibodies were purchased from Abcam. Rabbit anti-GFAP (Z0334) primary antibody was purchased from Dako. RNAscope probes Myocilin (506401-C3) and Mgp (463381) were purchased from Advanced Cell Diagnostics. Rabbit anti-IBA1 (PA5-21274), Lipofectamine™ 3000 Transfection Reagent (L3000001) and 8-well Chamber slide (177402) were purchased from Thermo Fisher. pAAV Rep/Cap 2/2 and adenoviral helper (pAdDeltaF6) plasmids were obtained from the Penn Vector Core (University of Pennsylvania). pCAG-GFP, pAAV-CMV-GFP-WPRE were gifts from the Cepko lab in Harvard Medical School. Ad5 adenovirus (Ad-CMV-GFP) was a gift from the Do lab in PolyU.

### Cell Culture

All cell culture media and supplies were obtained from Thermo Fisher. HEK293T, MEF, and 661W cells were cultured in Dulbecco’s modified Eagle’s medium (DMEM, high glucose) supplemented with 10% fetal bovine serum (FBS) and 1% penicillin/streptomycin (P/S). hTM cells were cultured in DMEM (low glucose) supplemented with 10% FBS and 1% P/S. ARPE cells were cultured in DMEM/F12 (50:50 mix) supplemented with 10% FBS and 1% P/S. All cells were maintained in 5% CO2 at 37°C.

### Animals

C57BL/6J (Strain # 000664) and *Rosa26^LSL-tdTomato^* (Strain # 007914) mice were purchased from The Jackson Laboratory. All mice were kept on a 12h light/12h dark cycle in the Laboratory Animal Research Unit, City University of Hong Kong. All animal procedures performed were approved by the Hong Kong Department of Health under Animals Ordinance Chapter 340, and animal care was carried out in accordance with institutional guidelines.

### LNP-mRNA formulation and characterization

The GFP-mRNA and SpCas9-mRNA were produced using MEGAScript T7 kit (Invitrogen, AM1334) with m1Ψ-5’-triphosphate (TriLink, N-1081) modification by in vitro transcription. Capping of the in vitro transcribed mRNs was conducted co-transcriptionally with the Anti Reverse Cap Analog (TriLink, N-7003). Poly(A) tails were added to the mRNA with Poly(A) Tailing Kit (Invitrogen, AM1350). The mRNA was further purified by LiCl precipitation as suggested by the T7 kit guideline. LNP-mRNA was formulated by microfluidic mixing. Briefly, ethanol solutions containing ionizable lipid, cholesterol, DSPC, and DMG-PEG2K, at molar ratios of 50:38.5:10:1.5, were mixed with 50 mM citrate buffer (pH 4.0) containing RNA in a microfluidic mixer by two syringe pumps at a volumetric ratio of 1:3. The total flow rate was 2 mL/min. A N:P ratio (amino lipid-to-oligonucleotide phosphate) of 5.6 between ionizable lipids and the nucleic acids was maintained throughout the study. LNP-mRNA was dialyzed twice using 20K MWCO dialysis devices (Thermo Fisher, 88405) in phosphate-buffered saline (PBS, pH 7.4) and concentrated with 100 kDa Amicon Ultra centrifugal filters (Millipore, UFC910024). All LNPs were stored at 4 °C until use. Size distribution, polydispersity index (PDI) and zeta-potential of LNPs were determined via dynamic light scattering using a Zetasizer Nano ZS (Malvern Instruments). mRNA encapsulation efficiency was determined using Quant-iT RiboGreen RNA reagent (Thermo Fisher, R11491) according to the manufacturer’s protocol.

### AAV production

pAAV-CMV-GFP-WPRE, Rep/Cap 2/2 and pAdDeltaF6 plasmids were mixed with polyethylenimine (PEI) and added to HEK293T cells. 24 hours after transfection, the cell medium was changed to DMEM only; 72 hours after transfection, supernatant and cells were collected separately. AAV2 in the supernatant was precipitated by PEG-8000 (8.5% wt/vol PEG-8000 and 0.4 M NaCl for 2hr at 4°C), then centrifuged at 7000 g for 20 min, and resuspended in virus lysis buffer (150 mM NaCl and 20 mM Tris, pH 8.0). AAV2 in the cells was released by three freeze/ thaw cycles and homogenizing in lysis buffer. The AAV2 resuspension was run on an iodixanol gradient, and viruses in the 40% fraction were collected. Recovered AAV virus particles were washed and condensed three times with cold PBS using 100 kDa Amicon Ultra centrifugal filters. Protein SDS-PAGE gels were run to determine virus titers.

### Cell internalization and image analysis

Approximately 10,000 cells were seeded per well of 96-well plates or 8-well Chamber slides and incubated with LNP-mRNA or Lipofectamine-pDNA, maintaining a 1 ng/μL nucleic acids concentration for 24 or 48 hours at 37°C. Cells in chamber slides were fixed in 4% paraformaldehyde (PFA) for 20 min and incubated with DAPI for 30 min before mounting. Cells were imaged using confocal microscope (Nikon A1HD25) under the same exposure parameters. Images were taken at a magnification of 20×. The fluorescence intensity and cell counting of images were analysed using Fiji ImageJ.

### *In vitro* cell viability

A cell density of 4,000 per well was plated in 96-well plates and allowed to grow to 60-70% confluency for 24hr at 37 °C. Then cells were treated with MC3-GFP, ALC0315-GFP or SM102-GFP respectively at a mRNA concentration of 1 ng/μL in the medium. All cells were incubated for an additional 24hr and MTT assay (Thermo Fisher, V13154) was performed to measure cell viability based on manufacturer’s protocol.

### Intravitreal injection

1- to 2-month-old mice were anesthetized with ketamine (100 mg/kg)/xylazine (10 mg/kg). Intravitreal injections were performed using a pulled angled glass pipette controlled by a FemtoJet (Eppendorf). The needle tip was inserted through the sclera at the equator, near the dorsal limbus of the eyeball, and entered the vitreous cavity. The injection dosages of each eye were 200, 100, 40 or 20 ng mRNA/injection for SM102-GFP, 100 ng total RNA/injection for SM102-SpCas9-sgtdT and SM102-SpCas9-sgMgp, 5×10^9 viral particles/injection for AAV2 and 2×10^7 viral particles/injection for Ad5 adenovirus. The injection volume was 1 μL for all vectors. 1% fluorescein solution was added for better observation.

### Intracameral injection

1- to 2-month-old mice old mice were anesthetized with ketamine (100 mg/kg)/xylazine (10 mg/kg). Intracameral injections were performed using a pulled angled glass pipette controlled by a FemtoJet (Eppendorf). The needle tip was inserted through the corneal margin and entered the anterior chamber without injuring lens or iris. The injection volume and dosage of each eye was 1 μL and 100 ng total RNA/injection for SM102-SpCas9-sgtdT. After delivery, the needle was left in place for 30s and withdrawn slowly to minimize leakage.

### Immunohistochemistry

The mice were sacrificed by CO_2_ euthanasia or cervical dislocation. The eyeballs were dorsally marked before enucleation, and were dissected into different tissues and fixed in 4% PFA for 30 min at room temperature. Fixed eyecups were washed three times with PBS and sequentially cryoprotected in 5%, 15%, and 30% sucrose for 15, 30, and 60 mins, respectively. Next, the eyecups were soaked in optimal cutting temperature embedding media and 30% sucrose solution at a ratio of 1:1 at 4℃ overnight and subsequently embedded in cryomold to achieve dorsal-ventral oriented slice. After tissue freezing below -20℃, a series of 20 μm sections were cut and collected in the glass slides using a cryostat machine (Thermo HM525NX Cryostat). During immunostaining, the retinal sections, fixed anterior segments or fixed retinas were first incubated in 3% BSA in PBST (PBS with 0.1% Triton X-100) for 1 hour, then were incubated with primary antibodies at recommended dilution at 4°C, overnight. After primary antibody incubation, the samples were washed three times with PBST before incubation in a mixture of DAPI (0.5 μg/ml) and secondary antibodies in the dark for at least 2 hours at room temperature. The retinal sections, anterior segment flat-mounts or retinal flat-mounts were washed again and mounted with an anti-fade solution. Slide images were captured using confocal microscope (Nikon A1HD25). All images were taken at a magnification of 20×. The image processing, histology measurements and cell counting were performed in Fiji ImageJ.

### *In situ* RNA hybridization

In situ RNA hybridization was performed using the RNAscope Multiplex Fluorescent Detection Reagents V2 kit (Advanced Cell Diagnostics) following the commercial guidelines. In general, eyecups were dissected, 4% PFA fixed, dehydrated in sucrose solution and embedded in OCT media. Eyecups were cryosectioned into 20 μm and mounted on SuperFrost Plus glass slides. After OCT removal by PBS and further dehydration by ethanol, retinal sections were stained with anti-Collagen-I antibody at 4°C overnight. After wash three times with PBST (PBS with 0.1% Tween-20), retinal sections were hybridized with Myocilin and Mgp probes for 2hr at 40 °C. Following the RNA hybridization steps, slides were stained using a mixture of DAPI and secondary antibodies for 2hr at room temperature. Slide images were captured using confocal microscope (Nikon A1HD25). All images were taken at a magnification of 20×. The image processing and histology measurements were performed in Fiji ImageJ.

### Tissue preparation and next generation sequencing (NGS)

SM102-SpCas9-sgMgp injected mouse eyes were harvested three weeks post-injection. The iridocorneal angle tissues, including TM strip, ciliary body, some cornea, and sclera, were dissected from the whole eyeball by removing the posterior segments, lens, iris, and most of the cornea. Genomic DNA was extracted from the remaining tissues using a PureLink™ Genomic DNA Mini Kit (Thermo Fisher, K182001) based on manufacturer’s protocol. A two-step PCR method, described in the previous study (*54*), was adopted to amplify the target sites of SpCas9-sg*Mgp*s from the extracted genomic DNAs. The libraries were sequenced on Illumina iSeq or NextSeq platforms and the sequencing results were analysed using CRISPResso2 (*55*).

### qPCR

Total RNA from retinas or TM tissues were extracted using TRIzol (Invitrogen) according to the manufacturer’s instructions. RNA purity and concentration were determined using Nanodrop spectrophotometry. RNA was converted to cDNA using a PrimeScript RT reagent kit with gDNAEraser (Takara Bio). qPCR was performed using the qPCR master mix (SybrGreen, Invitrogen) on QuantStudio 3 Real-Time PCR systems (Applied Biosystems). Target gene expression was normalized to mouse Gapdh.

### Intraocular Pressure (IOP)

An Icare TonoLab tonometer (Colonial Medical Supply) was used for noninvasive IOP measurement. Mice were anesthetized using 2.5% isoflurane, and IOP measurements were acquired from each eye within 3 min of induction of anesthesia. Each value was the average of six individual measurements. All measurements were performed at the same time during daylight (1:00-3:00 p.m.).

### Spectral-domain optical coherence tomography (SD-OCT)

OCT images of mouse eyes were taken using an SD-OCT (Bioptigen Envisu R4310 SD-OCT, Germany). Mice were anesthetized by intraperitoneal injection of a ketamine (100 mg/kg)/ xylazine (10 mg/kg) mixture at weight-adjusted dose. A drop of 0.5% tropicamide and 0.5% phenylephrine hydrochloride (Mydrin-P, Santen Pharmaceutical Co., Japan) solution was instilled on the ocular surface for pupil dilation for whole-eye biometry and retinal OCT measurement. Lubricating eye drops (Systane Ultra, Alcon) were applied to prevent desiccation of the cornea during imaging. Then, the anesthetized mouse was placed on a stereotaxic platform for alignment with the imaging lens. Whole-eye, retina and anterior chamber OCT images were separately measured and captured along the horizontal meridian centered at a point one optic disk diameter away from the outer optic disc margin using the SD-OCT. Axial resolution was 2.6 μm and scanning speed was 20,000 lines per second. Dimensions of individual ocular components were quantified using ImageJ. Axial length was defined as the distance between the anterior cornea and the outer boundary of the RPE layer.

### Electroretinography (ERG)

The eye physiology of mice was determined by ERG measurements using Espion E3 System (Diagnosys LLC). The mice were adapted in a dark cabinet for at least 3hr before ERG testing, then were anesthetized with a ketamine (100 mg/kg)/ xylazine (10 mg/kg) mixture. The eyes were kept hydrated with a topical gel. After putting mice on the platform, gold-wire electrodes for measuring electrical responses were placed on each eye cornea, while a reference electrode and a ground electrode were placed in the mouth and the tail, respectively. All these steps were performed in the darkroom under dim red light. For scotopic ERG recordings, 530nm light stimuli with different intensities (increments from 0.01 cd.s/m^2^ to 30 cd.s/m^2^) were elicited to stimulate scotopic responses in a specific time interval. For photopic ERG recordings, 5 min exposure under 10 cd.s/m^2^ light intensity was adopted to inhibit the rod function. The photopic response was measured by multiple flashes of 30 cd.s/m^2^ intensity in the illuminated background (10 cd.s/m^2^). The average amplitude and implicit time of a- and b-wave were recorded and exported for further analysis.

### Pattern-ERG (PERG)

The RGC function of mice eyes was determined by PERG measurements using Miami PERG system (Intelligent Hearing Systems, Miami, FL). Mice were anesthetized by intraperitoneal injection of a ketamine (100 mg/kg)/ xylazine (10 mg/kg) mixture at weight-adjusted dose, and then were placed on a heating pad to maintain animal core temperature at 37°C. The reference electrode was placed subcutaneously on the back of the head between the two ears and the ground electrode was placed at the root of the tail. The active steel needle electrode was placed subcutaneously on the snout for the simultaneous acquisition of left and right eye responses. Two 14 cm x 14 cm LED-based stimulators were placed in front so that the center of each screen was 10 cm from each eye. The pattern remained at a contrast of 85% and a luminance of 800 cd/m2, and consisted of four cycles of black-gray elements, with a spatial frequency of 0.052 c/d. Upon stimulation, the independent PERG signals were recorded from the snout and simultaneously by asynchronous binocular acquisition. With each trace recording up to 1020 ms, two consecutive recordings of 372 traces were averaged to achieve one readout. The first positive peak in the waveform was designated as P1 (typically around 100 ms) and the second negative peak as N2 (typically around 200 ms). The amplitude was measured from P1 to N2.

### Statistical Analysis

Data are presented as Mean ± SEM in all figures. Sample sizes and statistical analysis were indicated for each experiment in the figure legend. One-way ANOVA analysis followed by the Tukey test was performed to compare multiple groups, and Student’s t-test to compare two groups. A P value < 0.05 was considered statistically significant. GraphPad Prism was used to perform statistical analysis and create figures.

## Acknowledgements

This research was funded by the Innovation and Technology Fund from the Innovation and Technology Commission (GHP/223/21SZ), Hong Kong Research Grants Council Project (11103819, 11102922, and 11100723), Hong Kong Health and Medical Research Fund Project (05160276 and 06172466), TUNG Biomedical Sciences Foundation.

---

**Figure S1.**
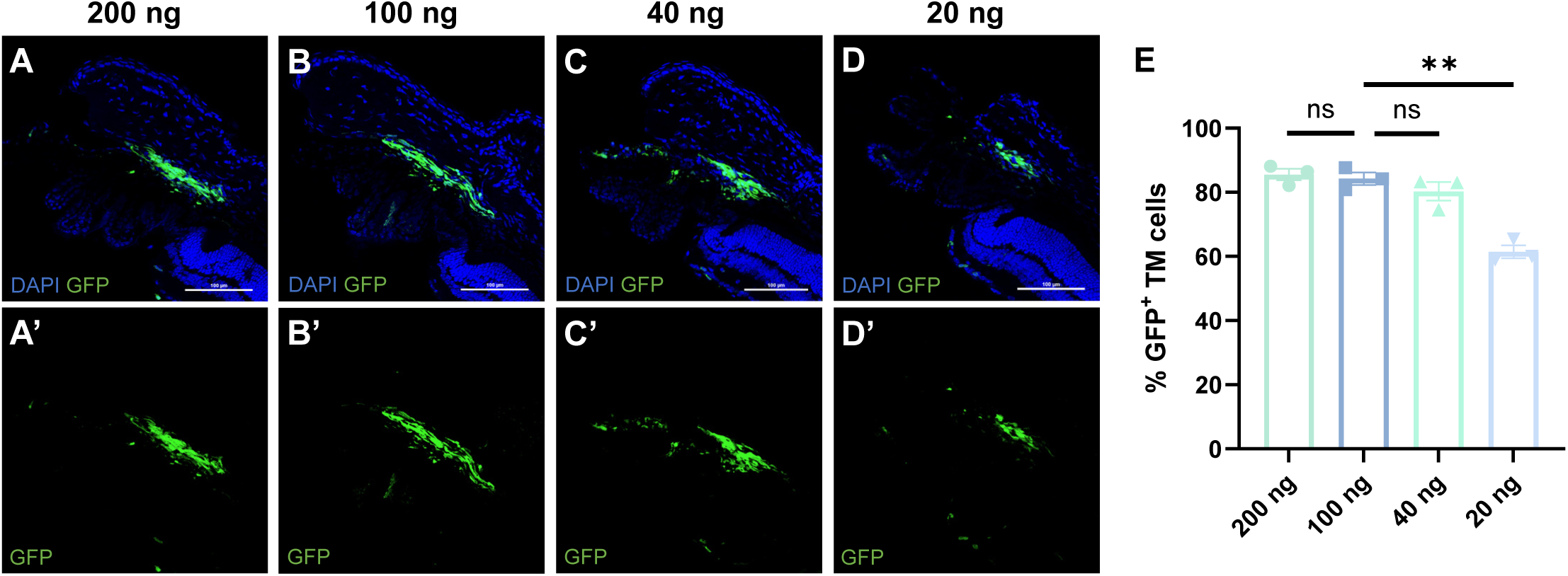
The dose study of SM102-GFP in TMs. (A-D) Representative images of retinal sections comparing TM infection effect by intravitreally-delivered SM102-GFP of different doses. (E) Quantification of GFP-positive TM cell numbers of eyes injected by SM102-GFP of different doses (n=3 for all groups). Data shown as Mean ± SEM. **P < 0.01; ns, not significant difference by one-way ANOVA analysis with Tukey test. Scale bars: 100 μm.

**Figure S2.**
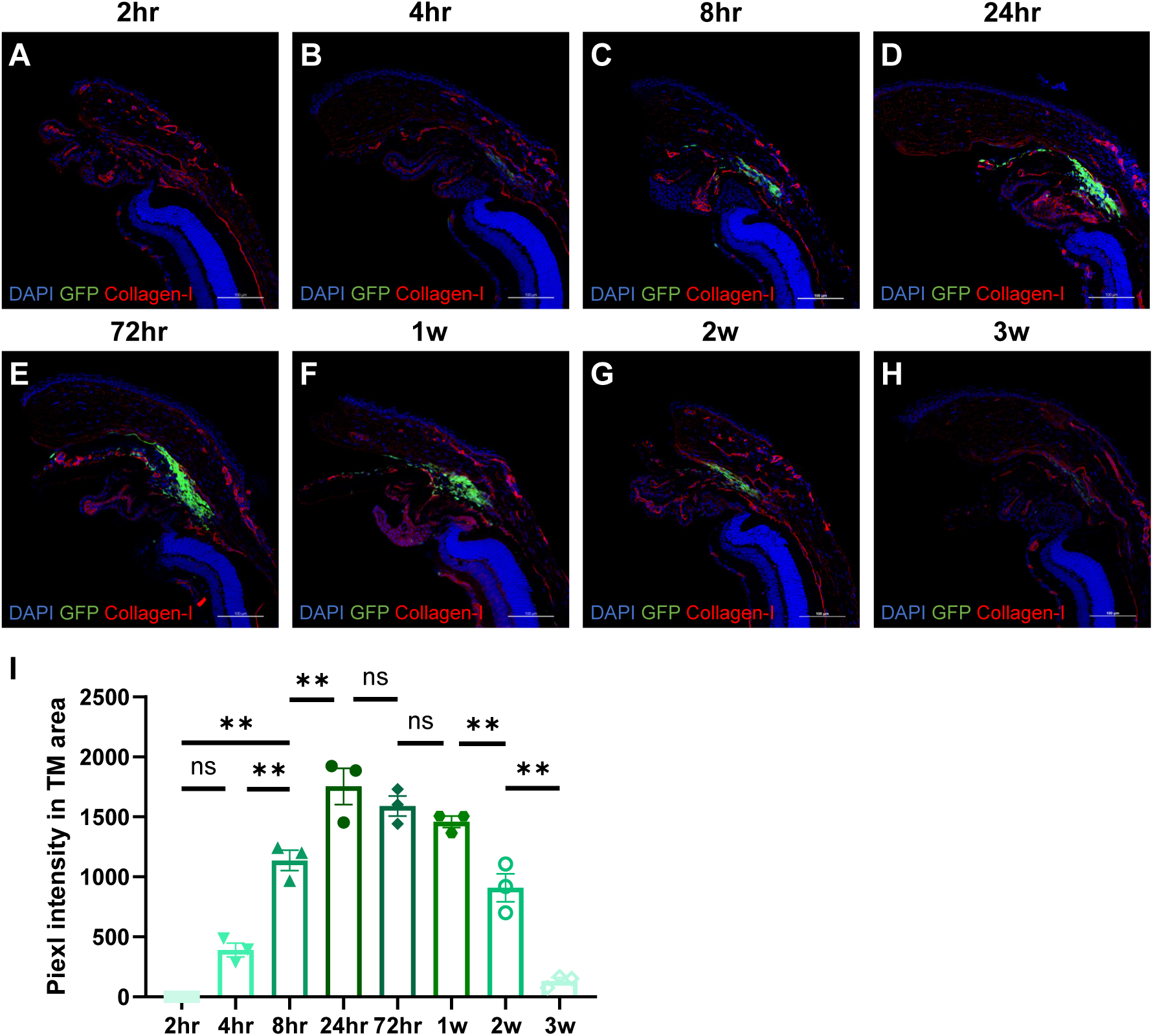
SM102-GFP enables rapid and transient mRNA expression to the mouse TM tissues. (A-H) GFP expression at different times-post injection showing the rapid and transient mRNA delivery feature of LNP in TMs by intravitreal injection. (I) Quantification of MFI in TM areas of SM102-GFP injected eyes harvested at different time courses (n=3 for all groups). Data shown as Mean ± SEM. **P < 0.01; ns, not significant difference by one-way ANOVA analysis with Tukey test. Scale bars: 100 μm.

**Figure S3.**
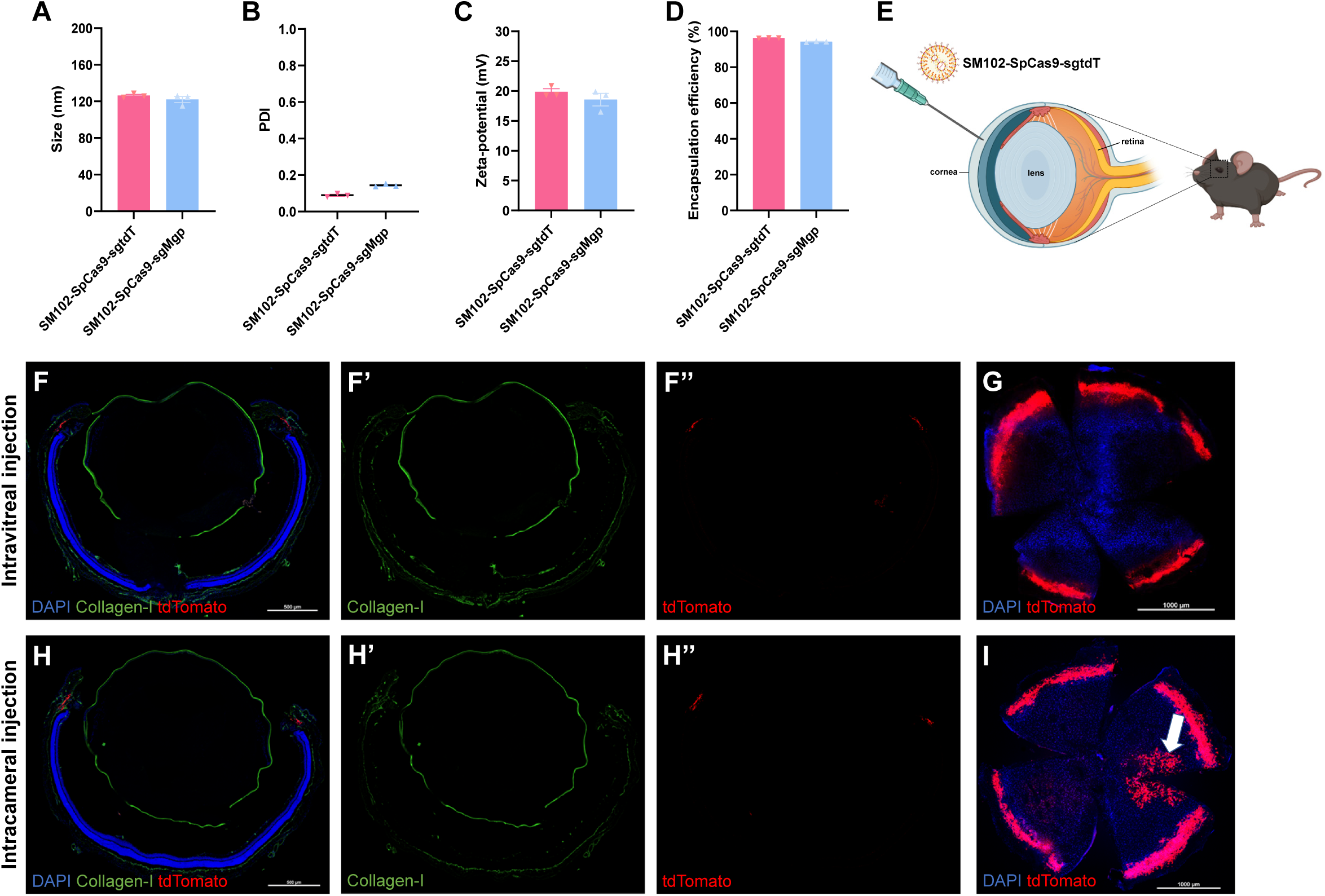
Both intravitreally- and intracamerally-delivered LNP-CRISPR mediates potent gene editing in TMs. (A-D) Characterizations of SM102-SpCas9-sgtdT and SM102-SpCas9-sgMgp, including size (A), polydispersity index (PDI) (B), zeta-potential (C), and encapsulation efficiency (D). (E) Schematic illustration showing intracameral delivery of SM102-SpCas9-sgtdT. (F-I) Representative images of retinal sections and anterior segment flat-mounts showing tdTomato expression in retinal outflow facility induced by intravitreally- or intracamerally-delivered SM102-SpCas9-sgtdT. The white arrow indicates cornea infection by intracamerally-injected SM102-SpCas9-sgtdT. Scale bars: 500 μm for whole retinal sections; 1000 μm for anterior segment flat-mounts.

**Figure S4.**
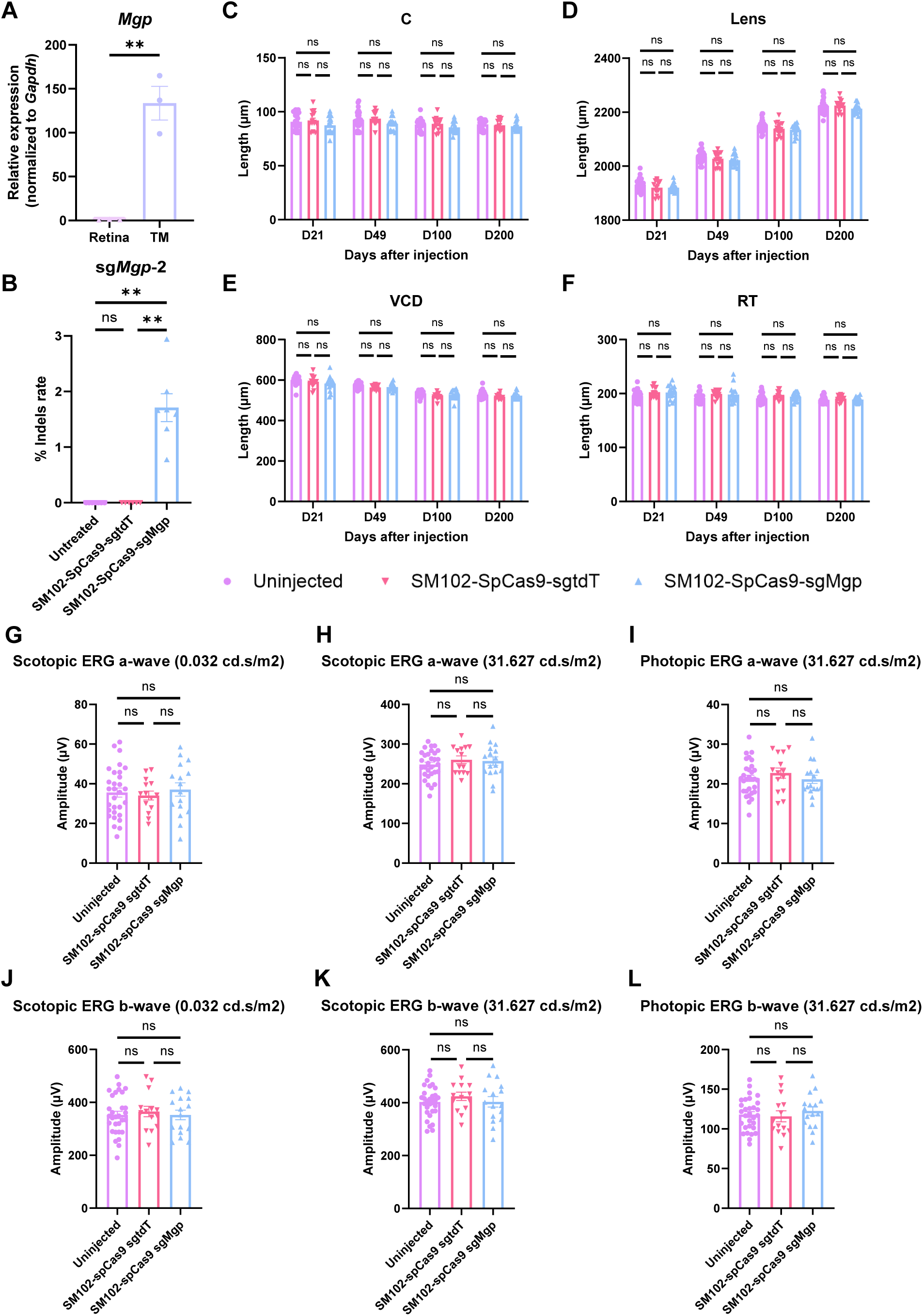
*Mgp*-KO eyes showed no variations in other ocular tissues and no retinal function impairment. (A) qPCR analysis of Mgp mRNA expression in mouse retina and TM tissues (n=3, normalized to Gapdh mRNA). (B) Quantification of insertions and deletions (Indels) at sg*Mgp*-2 targeting site in SM102-SpCas9-sgMgp (n=7) or SM102-SpCas9-sgtdT (n=6) treated or untreated (n=7) TM tissues by using next generation sequencing. (C-F) Quantification of the length of cornea (C), Lens, vitreous chamber depth (VCD) and retina (RT) of uninjected (n=30), control LNP-injected (n=14) and *Mgp*-KO eyes (n=16). (G-L) a- and b-wave amplitudes of scotopic and photopic ERG responses of uninjected (n=30), control LNP-injected (n=14) and *Mgp*-KO eyes (n=16) under light intensity 0.032 cd.s/m^2^ or 31.627 cd.s/m^2^ on Day 200-post injection. Data shown as Mean ± SEM. **P < 0.01; ns, not significant difference by two-tail unpaired student t-test (A) and one-way ANOVA analysis with Tukey test (B-L).

**Figure S5.**
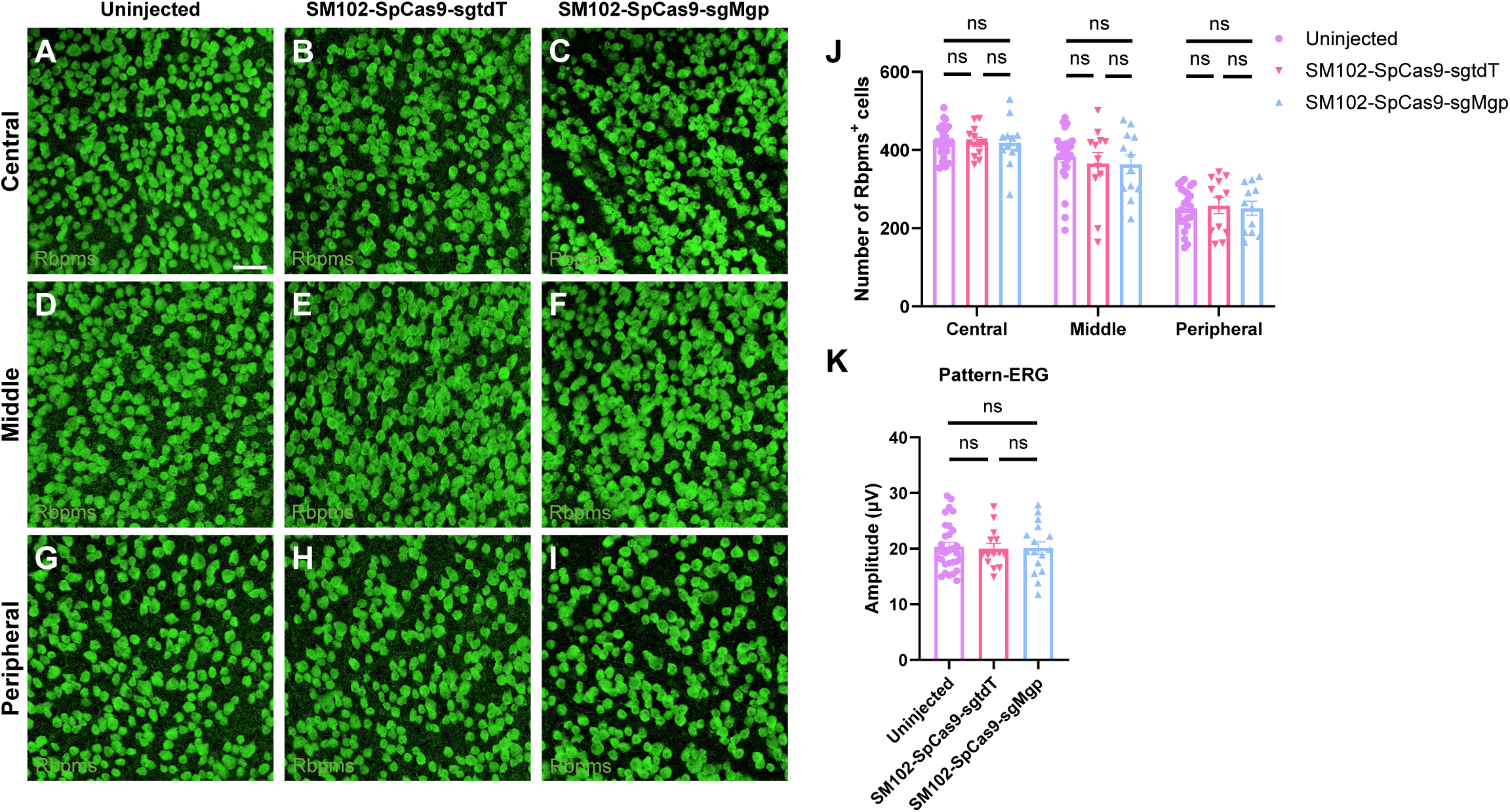
*Mgp*-KO eyes did not show massive RGC loss or attenuated RGC function. (A-I) Representative zoom-in images of retinal flat-mounts harvested on Day 200-post injection with anti-Rbpms staining showing the number of RGCs at the central, middle and peripheral retinal positions. (J) Quantification of Rbpms-positive RGC numbers of uninjected (n=24), control LNP-injected (n=12) and *Mgp*-KO eyes (n=12). (K) P1-N2 amplitudes of pattern-ERG responses of uninjected (n=30), control LNP-injected (n=14) and *Mgp*-KO eyes (n=16) on Day 200-post injection. Data shown as Mean ± SEM. ns, not significant difference by one-way ANOVA analysis with Tukey test. Scale bars: 50 μm.

